# Fitness effects of adult crowding in *Drosophila*: more than just overall density

**DOI:** 10.1101/2025.04.24.650416

**Authors:** Medha Rao, Chinmay Temura, R. S. Bindya, Amitabh Joshi

## Abstract

Density-dependent selection has been widely studied in *D. melanogaster* in the context of larval crowding, revealing its impact on several life-history traits. However, the effects of adult crowding in *Drosophila* have not been studied in similar detail. Earlier studies on adult crowding have primarily used large flies (reared at low larval density). These gave rise to the notion that adult crowding negatively affects key fitness components such as mortality and fecundity. Earlier work from our lab showed that body size significantly alters how flies respond to a relatively short period of adult crowding. Large flies show increased mortality and decreased fecundity with increased adult density. In contrast, small flies (reared at high larval density) tolerate adult crowding better, showing increased fecundity with increased adult density. Here, we extend this line of work by investigating how air volume in a culture vial can influence the effects of adult crowding. We manipulated air volume by altering the vial diameter (thereby also the food surface area), or the height of the air column. Flies of different body sizes were generated by rearing them at low or high larval densities. We find that the surface area of food available to the adults plays a greater role in shaping the outcome of adult crowding than the height of the air column. Large flies displayed context-dependent responses to adult crowding that were driven by the surface area of food provided. Small flies consistently responded positively to adult crowding in all conditions, with low mortality and increased fecundity at high adult density. Our findings highlight the importance of considering the body size, absolute density and the food surface area available to the flies and their nuanced interactions while exploring the effects of adult crowding, and thus paves the way for a detailed examination of other factors that impact adult crowding in addition to just the overall density.

## Introduction

Density-dependent selection in *D. melanogaster* has been extensively studied in the context of larval crowding, revealing its impact on various life-history related traits such as larval feeding rates (Joshi & Mueller, 1988, 1996), waste tolerance (Shiotsugu *et al*., 1997; Borash *et al*., 1998), pupation height (Joshi & Mueller, 1993, 1996), egg-to-adult development time and egg-to-adult survivorship (reviewed in Mueller, 1997). However, the effects of adult crowding in *Drosophila* have not been studied in as much detail (reviewed in Prasad & Joshi, 2003). Studies examining adult crowding in *Drosophila* have mostly been single-generation experiments (Pearl & Parker, 1922; Pearl, 1927; Pearl *et al*., 1927; Pearl, 1932; Robertson & Sang, 1944), while a few studies used an experimental evolution approach to examine the effects of adult crowding (Joshi, 1997; Joshi *et al*., 1998; Borash & Ho, 2001). The results from these studies have led to the formation of a widely held notion that adult crowding has a negative effect on fitness components such as mortality and fecundity. However, across these studies, the effects of adult crowding were tested using only large flies obtained by rearing flies at low larval density (L-LD). As the space available during crowding is limited, body size could affect the response of flies to episodes of adult crowding. Thus, smaller flies might experience the harmful effects of adult crowding to a lower degree than large flies.

In an earlier report, we tested the validity of the notion that the number of individuals localized in a fixed space could be used as a reliable index of the intensity of adult crowding (Rao *et al*., 2025). We showed that the body size of the flies significantly influenced the net effect on mortality during, and female fecundity after, an episode of adult crowding. Large flies (reared at L-LD) showed a response typically associated with adult crowding – an increase in mortality and a pattern of decreasing fecundity when subjected to adult crowding (Pearl, 1932; Robertson & Sang, 1944; Joshi *et al*., 1998). In contrast, small flies, obtained by rearing at high larval density (H-LD) or by enforcing selection for rapid development, tolerated adult crowding better, exhibiting low mortality and increased fecundity with adult density. The results highlight the importance of considering both body size and density while studying responses to adult crowding.

Parallels can be drawn to studies that examined larval crowding in *Drosophila,* highlighting the importance of a nuanced approach to studying how larval crowding was imposed and their subsequent impact on the traits that evolved in populations (Nagarajan *et al*., 2016; Sarangi, 2018; Venkitachalam, 2023). These studies revealed that subtle variations in larval rearing conditions – including egg number, food volume, and surface area of the food column – critically shape life-history traits like pre-adult development time, pre-adult survivorship and adult biomass at eclosion (Nagarajan *et al*., 2016; Sarangi, 2018; Venkitachalam, 2023). It was also found that *effective larval density* (number of larvae present in the feeding band) was a better predictor of the outcome of larval crowding than the *overall density* (number of larvae present in the total volume of food; Venkitachalam *et al*., 2023). These results emphasized that differences in the ecological details of how larval crowding was implemented could significantly affect the outcome of crowding on the various fitness-related traits mentioned above and highlight the importance of a nuanced approach while studying the effects of larval crowding. However, studies addressing similar nuances in adult crowding are quite rare.

Earlier work on adult crowding revealed a pattern of higher mortality and lower fecundity at higher adult densities (Pearl, 1932; Robertson & Sang, 1944; Joshi *et al*., 1998). The high mortality across these studies was believed to be caused by the sogginess of the medium (due to increased larval activity), leading to increased female mortality as they visit the food surface more often than males for feeding and oviposition (Joshi, 1997; Joshi *et al*., 1998). One study defined the density of a population as the number of individuals per unit of volume or area of the universe concerned (Pearl, 1927). Based on this definition, their experimental studies focused on altering adult density primarily by changing the number of individuals. In a subsequent study (Pearl, 1932), the author aimed to change the adult density by modifying the air volume available to flies during crowding. In this experiment, while the air volume was different across treatments (half-pint and quarter-pint bottles), the surface area of food available was uniform. Fecundity and mortality were similar across both treatments, indicating that air volume did not have a significant effect. The author speculated that the surface area of food available might play a critical role in shaping the intensity of crowding, an idea that was not tested subsequently.

Another study aiming to replicate Pearl’s results found that the absolute values of female fecundity were much higher in their study (Robertson & Sang, 1944). In this case, the containers used to house flies (dipping jars) had an air volume similar to half-pint bottles, but the surface area of food provided was much higher (∼77.4% larger). The authors postulated that the higher fecundity observed was either due to the strain of flies used in the assay or due to the differences in the surface area of food available to the adults. Although these studies inferred a prominent role of the surface area of food in shaping the effects of adult crowding, that conjecture is yet to be systematically tested. For instance, the differences in fecundity observed across the studies could also have been due to random variation across assays. Additionally, while the containers had an equal air volume across studies, they were of different dimensions. Thus, apart from the surface area of food, the height of the air column could also potentially contribute to the observed differences. However, these aspects were not revisited formally again, and it would be informative to systematically demonstrate the effects of changes in the parameters, as mentioned earlier, using better experimental designs.

In this study, we aimed to extend this line of work by examining the effects of altering the air volume available to adults during an episode of adult crowding on mortality and female fecundity (Fig. 1). The air volume was altered by modifying either the air column height or the vial diameter. Modifying the vial diameter would also alter the surface area of the food available to the adults for feeding/oviposition and provide evidence of the impact of the surface area of food during adult crowding.

**Figure 1:**
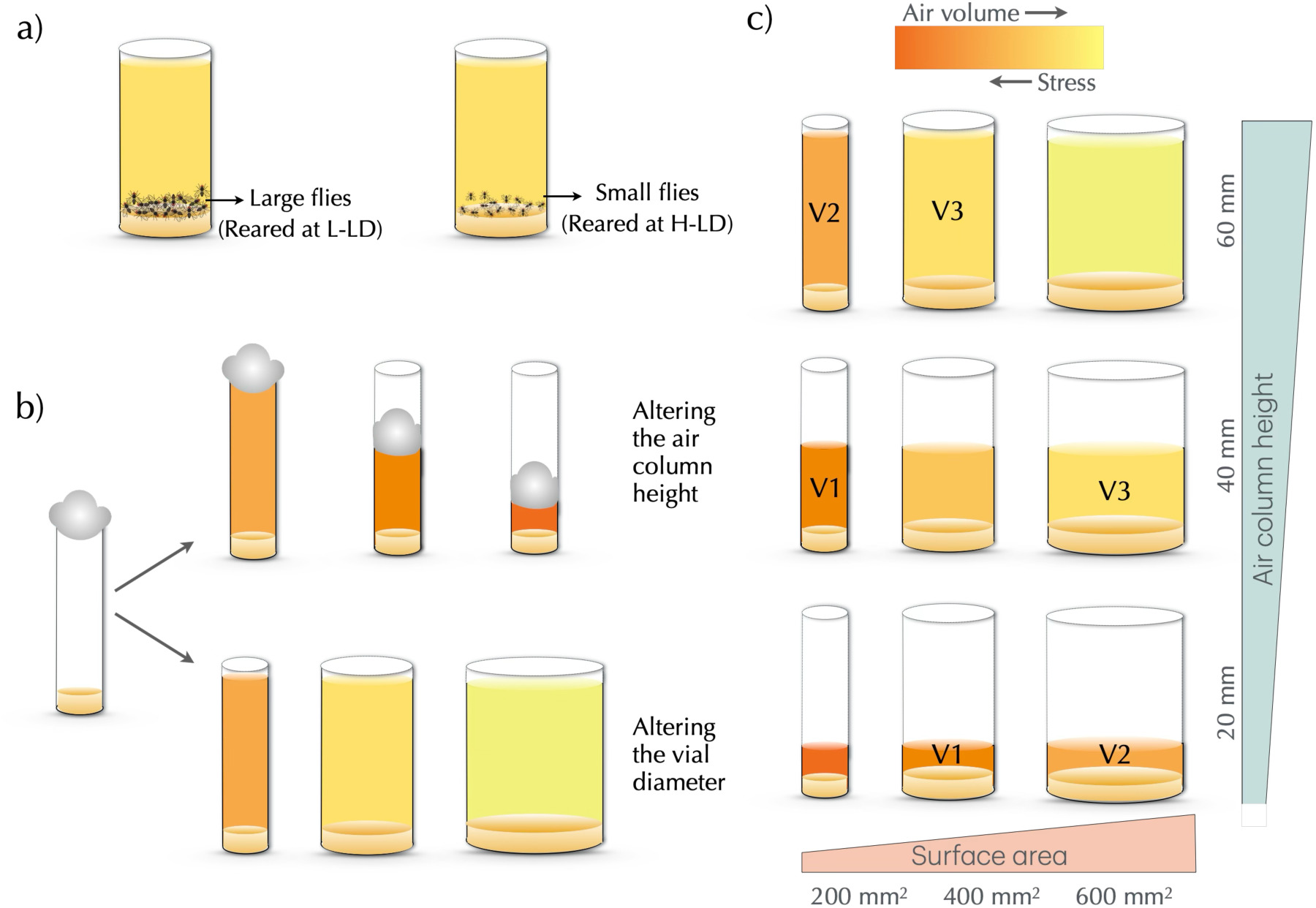
The experimental design. **a)** Flies of varying body sizes can experience different levels of stress, even when the number of flies per vial and the available air volume is equal. **b)** Air volume available to the flies can be altered in one of two ways – by altering the air column height (changing the position of the cotton plug within the vial) or by altering the vial diameter (also resulting in a change in the available food surface area). **c)** Nine treatment combinations are generated through changes in vial dimensions based on air column height or vial diameter. Three pairs of treatment combinations with equal air volume are marked V1-3. Flies reared at either low or high larval density underwent adult conditioning (either low or high adult density) for two days in these vials resulting in a total of 36 treatment combinations per replicate population.

Building on our earlier findings, we had postulated that the stress experienced by large flies at high adult densities (H-AD) would be significantly higher than that experienced by small flies (Rao *et al*., 2025). The stress experienced during crowding could increase mortality and lower the fecundity of the surviving females. We hypothesized a hump-shaped relationship between female fecundity and adult density, suggesting that this relationship was affected by the body size of the individuals. For large flies, fecundity initially increases with density (zone of positive effects) and subsequently decreases (zone of adverse effects), whereas, for small flies, this threshold may occur at much higher adult densities, resulting in a larger zone of positive effects (see Figure 4 in Rao *et al*., 2025). Therefore, the *effective adult density* within the container - reflecting the number of individuals, their body size (total biomass) and the air volume (or perhaps the food surface area) – should be the principal index of the net effect of adult crowding on fitness components.

Our present findings reveal that vial diameter had a more prominent effect on mortality and female fecundity than the air column height. Large flies showed a context-dependent response to adult crowding, primarily driven by the surface area of the food provided to the adults. In contrast, small flies, obtained here by the imposition of larval crowding, consistently showed a pattern of increased fecundity upon imposition of H-AD under all treatment combinations. Therefore, the effect of changes in surface area or air column height on the response to adult crowding was contingent on the body size of the flies.

## Materials and Methods

### Study populations

Four large (∼1800 individuals), laboratory-maintained, replicate *D. melanogaster* populations (Joshi <u>B</u>aseline; JB_1-4_) were used for the study. Their maintenance protocols and ancestry have been reported in detail earlier (Sheeba *et al*., 1998) and will be summarized below. At the time of the assay, JBs had undergone at least 457 generations of maintenance in the laboratory.

### Maintenance protocol

The JBs are maintained at 25±1°C, high humidity and constant light on banana-jaggery medium. Eggs are collected in 8-dram vials (2.2 cm dia × 9.6 cm ht) at a density of ∼70 eggs in ∼6 mL of banana-jaggery medium. Once all the adults emerge (day 12 from egg collection), they are transferred to adult collection vials containing ∼4 mL of food. The adults undergo two more rounds of vial-to-vial transfers on days 14 and 16 to receive fresh food. On day 18, the flies are transferred to a Plexiglas cage (25 × 20 × 15 cm^3^) and provided with a Petri plate containing food medium coated with yeast paste (henceforth, yeasted food plate) for two days. Following this, a fresh food plate cut into two halves (henceforth, egg-laying plate) is provided for oviposition. The vertical edges created by cutting the food stimulate oviposition in the females. After 18 hours, eggs from this plate are collected to initiate the next generation. Thus, the JBs are maintained on a 21-day discrete generation cycle.

### Standardization of populations

Before the assay, we imposed one generation of common rearing conditions for all populations. Eggs were collected from each replicate population at a density of ∼70 eggs in 6 mL of food. Once the eclosion of adults was completed (day 12 from egg collection), all the adults were directly transferred to a Plexiglas cage, and a yeasted food plate was provided. After two days, the flies were provided with an egg-laying plate for 18 hours. The eggs collected from these plates were used for the assay egg collection.

### Assay egg collection

The assay egg collection involved initiating two types of larval density cultures from each replicate population – a low larval density (70±10 eggs in 6 mL: L-LD) and a high larval density (300±20 eggs in 2 mL: H-LD) culture. The egg collection day is denoted as Day 0. The assay was performed on one replicate population (block) at a time for logistic reasons.

### Collection of flies

All eclosed flies from the L-LD treatment were transferred to Plexiglas cages on day 12 from egg collection. In H-LD cultures, the eclosion of the adults spanned over several days (between days 9 and 16), and eclosed adults from these cultures were transferred to Plexiglas cages once daily during this time period. All cages were provided with fresh food on alternate days.

### Adult crowding

Flies were exposed to one of two adult conditioning densities for two days *before* measuring female fecundity (between days 16 and 18). The conditioning treatments were set up on day 16 from the egg collection. The low adult density (L-AD) treatment had five males and five females per vial, and the high adult density (H-AD) treatment had 75 males and 75 females per vial. This setup was carried out by anaesthetizing flies using CO_2_, and care was taken to ensure that flies spent minimal time on the CO_2_ pad. Ten replicate vials of L-AD and four replicate vials of H-AD were set up for each combination of larval density, vial diameter, air column height and replicate population.

The volume of air available to the adults was altered during the adult conditioning period in the following ways –

1. Change in the volume via *a change in the height of the air column* (height of the cotton plug from the food surface). The three heights selected were 20 mm, 40 mm and 60 mm from the surface of the food. Of the three, the 60 mm treatment represents the usual plug height from the food surface during routine population maintenance.
2. Change in volume via *a change in the vial’s cross-sectional area (CSA)*. We used three types of vials with CSA: 200 mm^2^, 400 mm^2^ and 600 mm^2^, which are hereafter referred to as narrow, regular and wide vials respectively. The 400 mm^2^ treatment represents the vial diameter used in routine population maintenance.

To ensure flies were provided the same range of heights across all three vial types during adult conditioning, we chose to keep the height of the food column in the adult collection vials uniform across the different vial types (set to 10 mm). Consequently, the volumes of food medium present vary across the three vial types, although the food available for flies was ad-libitum in all cases. All vials also had ad-libitum yeast paste applied onto one side of the vial wall. Ten replicate vials of L-AD and four replicate vials of H-AD were set up for each combination of larval density, vial diameter, air column height and replicate population.

Sex-specific mortality in the conditioning vials was recorded on day 18. The surviving females were aspirated and transferred to fresh vials containing ∼0.5 mL of banana-jaggery medium (to facilitate accurate egg counts). They were given an oviposition window of 24 hours, after which the number of eggs in each vial was counted. Twenty replicate vials were set up for each combination of larval density, adult density, surface area, column height and replicate population.

The present study differed in experimental design from our earlier report (Rao *et al*., 2025) in the following ways –

a) During the oviposition window for assaying fecundity, a single female was present instead of a mating pair. This was done to prevent any effect the presence of a male might have on the fecundity of the female during the oviposition window.
b) Flies were anaesthetized using CO_2_ to set up the adult conditioning vials for logistic reasons. Extreme care was taken to minimize the exposure of adults to the CO_2_.

The experiment involved a full-factorial design involving four fixed factors (larval and adult density, air column height, vial diameter) and a random factor (blocks; subsumes replicate populations and accounts for variation in replicate runs of the assay) crossed with each other. This design also provided us with three individual sets of the same air volume, achieved through different combinations of column height and vial diameter (Fig. 1c).

### Dry weight measurements

On the day of the adult conditioning set up (day 16 from egg collection), a set of flies from each combination of larval density and replicate population were collected from the Plexiglas cages for dry weight measurements. Thus, the collected flies represented the weight of the flies just before experiencing the adult conditioning treatment. The flies were transferred to empty vials, frozen and stored at -20℃. All flies were sorted by sex into five replicate batches of five flies each and dried in a hot air oven at 70°C for 36 hours. Triplicate weight measurements were obtained for each batch on a Sartorius (CP 225D) fine balance, and the average of these readings was considered as the weight of the batch.

### Statistical analyses

Mortality, fecundity and the dry weight data were subjected to a full-factorial mixed model Analyses of Variance (ANOVAs) at a significance level of 0.05. The cell mean of each treatment combination was the unit of analysis. Larval density (two levels), adult density (two levels), vial diameter (three levels) and air column height (three levels) were considered as fixed factors in the analysis for female fecundity. For the adult mortality ANOVA, sex of the fly (two levels) was a fixed factor in addition to the above-mentioned factors. The ANOVA of dry weights involved only larval density and sex as the fixed factors. In all the analyses, replicate populations (blocks) were treated as a random factor (four levels) and all the factors (fixed and random) were crossed with each other. Pair-wise post-hoc comparisons were performed using Tukey’s HSD at a significance level of 0.05.

Linear least-squares regressions were performed to determine if the surface area of food or the air volume better predicted the effect of adult crowding on mortality and fecundity. In each case, the number of adults per unit surface area or the number of adults per unit volume was used as the predictor variable, while mortality incurred during conditioning and female fecundity were used as response variables, respectively.

All ANOVAs were performed using R release 4.3.1 (R Core Team, 2024) using the *aov* function in the *stats* package (R Core Team, 2024). Regression analyses were performed in R using the *lm* function in the *stats* package (R Core Team, 2024). Graphs were plotted using R packages *tidyverse* and *ggplot2* (Wickham, 2016; Wickham et al., 2019).

## Results

We note that the results presented below are relevant to the significant effects found in the ANOVA (Tables S1-S3), and the figures for the full experimental design can be found in the supplementary materials (Fig. S1 & S2).

### An increase in food surface area reduces mortality and increases fecundity

Vial diameter significantly affected adult mortality and female fecundity. On an average, the highest mortality was observed in the narrow vials, followed by the regular and wide vials, respectively (Fig. 2a, Table S1, main effect of vial diameter, *F*_2,6_ = 153.89, *P* < 0.001). However, this effect was limited to flies exposed to H-AD. There was no difference in mortality at low adult density (L-AD) across the vial types (Fig. 2a, Table S1, interaction effect of vial diameter and adult density, *F*_2,6_ = 135.16, *P* < 0.001).

**Figure 2:**
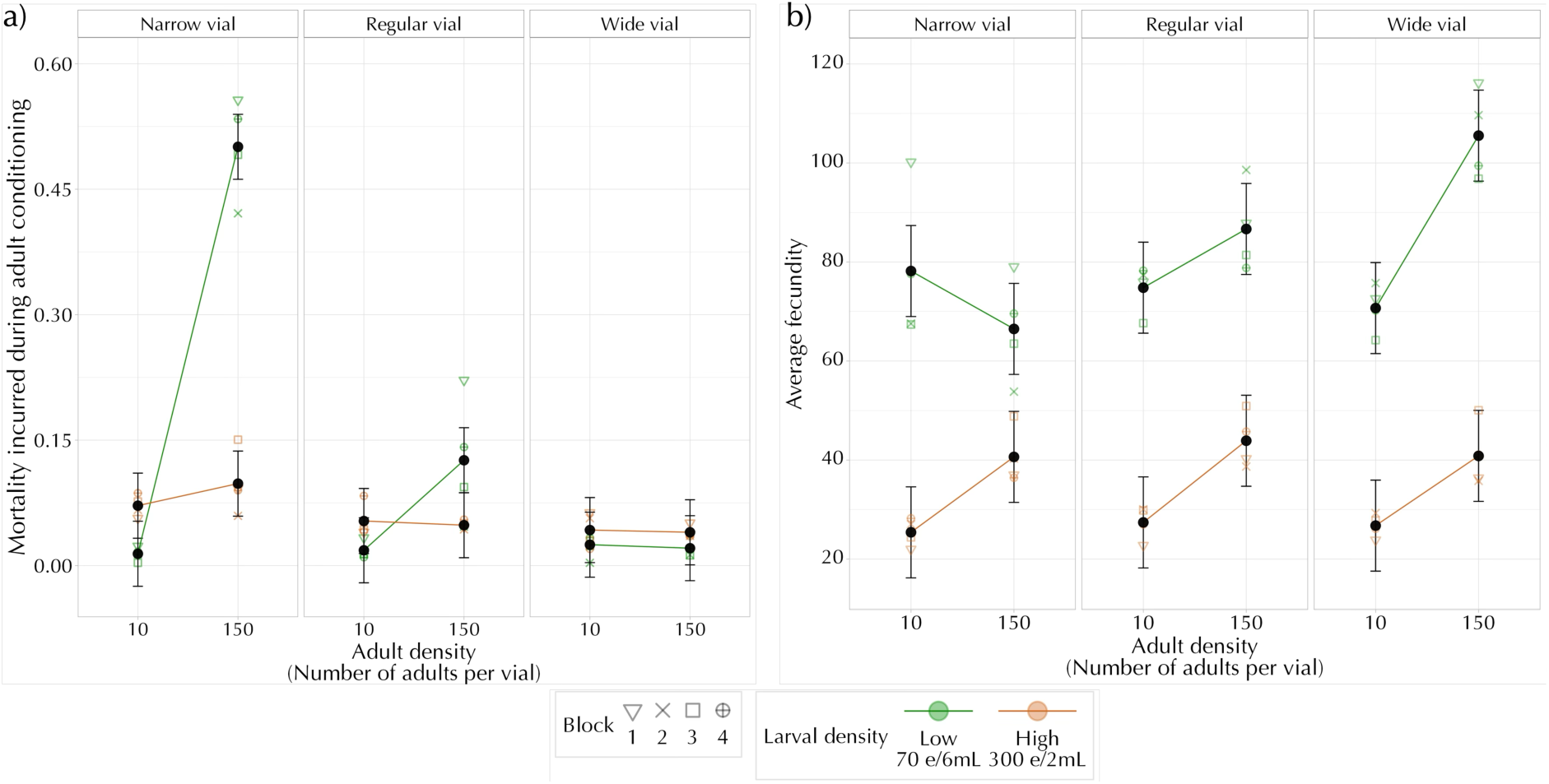
Effect of vial diameter, larval and adult density (averaged over other factors) on **a)** mean mortality during the adult conditioning and **b)** mean female fecundity measured after the conditioning period. Error bars are 95% confidence intervals around the mean and can be used for visual hypothesis testing.

The fecundity of the females was the highest in the case of wide vials, followed by regular and narrow vials, respectively (Fig. 2b, Table S2, main effect of vial diameter, *F*_2,6_ = 5.29, *P* = 0.047). As in the case of mortality, the impact of vial diameter was only observed at H-AD, and no significant differences were observed at L-AD (Fig. 2b, Table S2, interaction effect of vial diameter and adult density, *F*_2,6_ = 42.43, *P* < 0.001).

### The response of large flies to adult crowding depends on the vial diameter

Flies reared at L-LD developed into heavier adults than those reared at H-LD (Fig. S3). Female flies were significantly heavier than male flies across both larval rearing densities, with the difference between their weights being higher when reared at L-LD (Fig. S3, Table S3). When compared to small flies (reared at H-LD), large flies (reared at L-LD) experienced significantly higher mortality at H-AD (Fig. 2a, Table S1, interaction effect of larval and adult density, *F*_1,3_ = 101.74, *P* = 0.002). However, both large and small flies showed an increase in fecundity with an increase in adult density when averaged across various air volume treatments (lack of an interaction effect between larval and adult density; Fig. 2b, Table S2).

Large flies also showed a context-dependent response to adult crowding based on the vial diameter. In regular and narrow vials, high adult densities induced higher mortality. A considerable increase in mortality (∼45%) was observed when large flies were subjected to crowding in the narrow vials (Fig. 2a). There was no significant effect of increasing adult density on mortality in the wide vials. In the wide vials, a considerable increase in fecundity with an increase in adult density was observed, whereas, in the regular and narrow vials, there were patterns of non-significant increase and decrease in fecundity, respectively (Fig. 2b).

### Small flies showed a consistent response to changes in adult density or vial diameter

For small flies, there was no significant effect of adult density on mortality in any of the vial types (Fig. 2a). However, these flies showed a consistent pattern of increase in fecundity with an increase in adult density regardless of the vial diameter, which was strikingly similar to the response shown by the small flies in an earlier study (see Fig. 3 in Rao *et al*., 2025). The extent of the increase in fecundity was also very similar across the different vial types (Fig. 2b).

**Figure 3:**
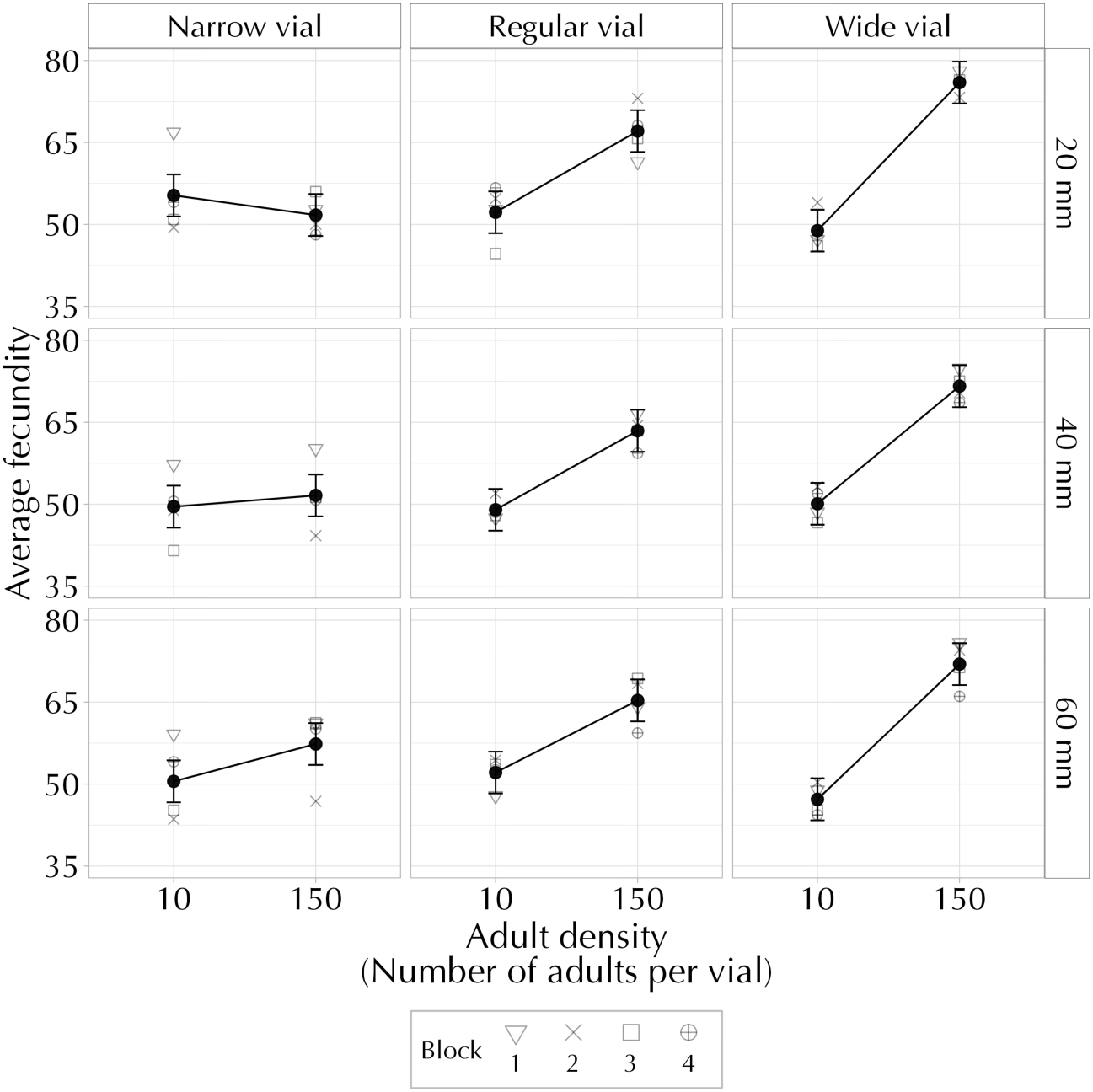
Effect of vial diameter, height of the air column and adult density on mean female fecundity measured after the conditioning period (averaged over other factors). Error bars are 95% confidence intervals around the mean and can be used for visual hypothesis testing.

### The height of the air column did not have a large effect on female fecundity

The height of the air column did not play any major role in shaping the effects of adult crowding on mortality, as evident from the lack of significant main or interactive effects (Table S1). Female fecundity was affected by the height of the air column (Fig. 3, Table S2, main effect of column height, *F*_2,6_ = 5.743, *P* = 0.04), with fecundity being significantly higher in the 20 mm treatment compared to the 40 mm treatment. However, we believe this is not of biological significance, as it amounted to an absolute difference of approximately two eggs (∼4.6% difference) between the treatments.

The patterns of fecundity were not altered to a great extent when the change in air volume was brought about through a change in the height of the air column, while the vial diameter was constant (Fig. 3). In comparison, a change in air volume brought about via a change in vial diameter alone resulted in significant changes in fecundity patterns. While on the one hand, a significant main effect of the height of the air column on fecundity was observed, our analyses indicate that in comparison to the vial diameter, the height of the air column had a relatively minor role to play in determining effects of adult crowding on female fecundity. It is further evidenced by the absolute difference of just two eggs between the 20 mm and 40 mm height treatments.

The experimental design included three equal-volume combinations, achieved through combinations of various vial diameters and air column heights (Fig. 1c). If the air volume played a prominent role in determining the intensity of crowding stress, we would expect to see a similar response to adult crowding in containers with equal air volume. However, the response shown by the flies was different across containers with equal volumes (Fig. 4a-c). The fecundity patterns seemed affected primarily by the vial diameter, as observed earlier (Fig. 3). In each set of comparisons at H-AD, the counterpart with the larger vial diameter showed lower mortality and higher fecundity than the counterpart with the smaller vial diameter, even though air volumes were identical. Thus, the surface area of food available to the adults during crowding appears to be more instrumental in determining the response to adult crowding rather than the volume of air provided.

**Figure 4:**
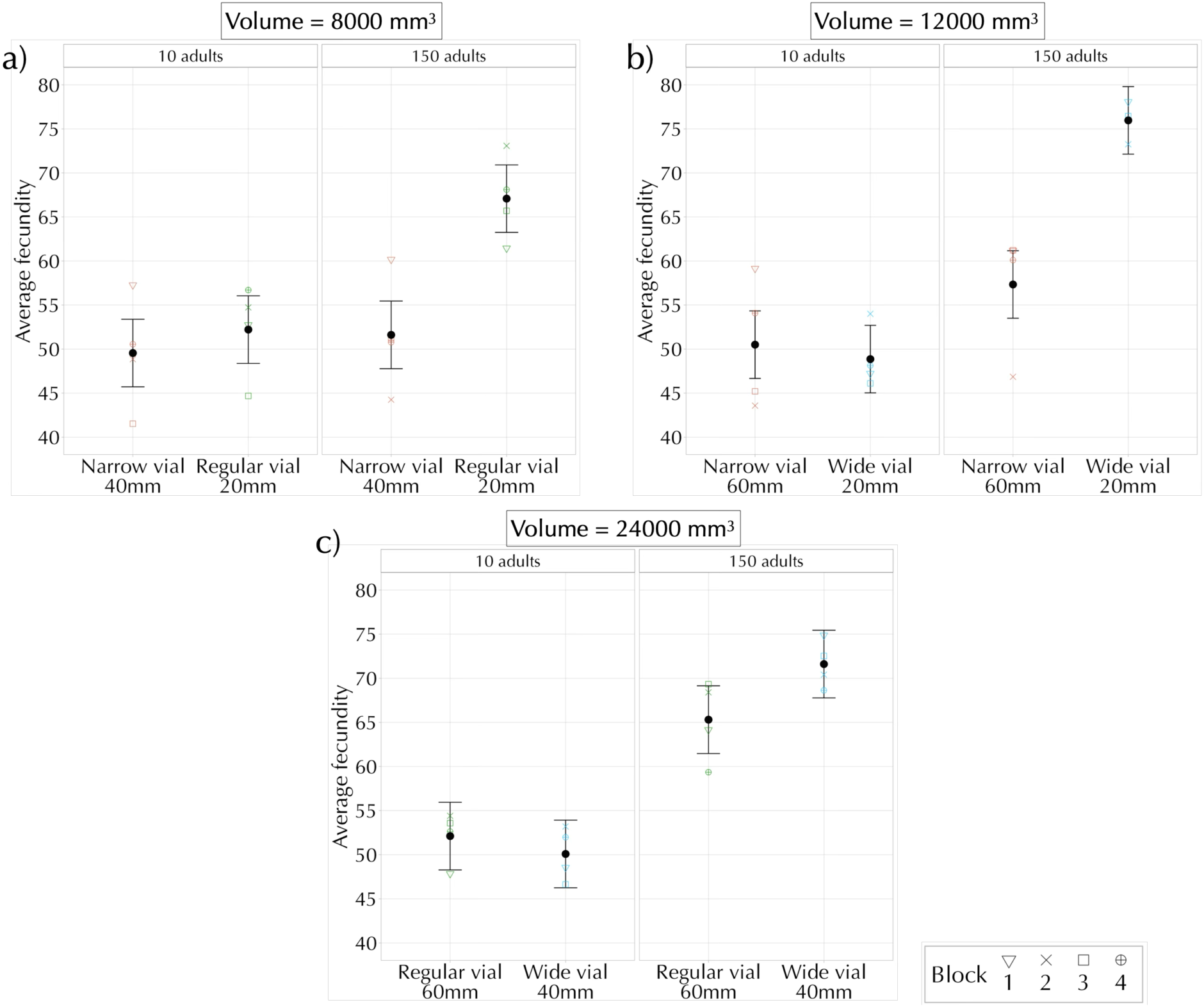
Effect of vial diameter, height of the air column and adult density on mean female fecundity measured after the adult conditioning period (averaged over other factors). Combinations depicted in **a)**-**c)** have an equal air volume available to the adults, achieved through different container dimensions (see Figure 1). Error bars represent 95% confidence intervals around the mean and can be used for visual hypothesis testing.

### For large flies crowded as adults, the food surface area is a better predictor of mortality and fecundity than the air volume

When using the entire dataset for regressions, the *R*^2^ values for mortality or fecundity were relatively low for either the number of adults per unit surface area (hereafter referred to as surface area) or the number of adults per unit air volume (hereafter referred to as air volume) as a predictor variable (Table S4). This finding is unsurprising as only the large flies responded to any changes in the surface area of food available (Fig. 2). Furthermore, these effects were observed only at H-AD, with negligible effects observed at L-AD. Based on this, regressions were carried out on subsets of the original data by using combinations of the level of larval and adult density experienced. These results are elaborated below, while the summary for the remaining regressions can be obtained from Table S4. For the large flies at H-AD, the surface area was a better predictor of adult mortality (*R*^2^ = 0.922, *P* < 0.001; Fig. 5a) than the air volume (*R*^2^ = 0.445, *P* < 0.001; Fig. 5b). Similarly, the surface area was a better predictor of female fecundity (*R*^2^ = 0.674, *P* < 0.001; Fig. 5c) compared to the volume of air available during crowding (*R*^2^ = 0.319, *P* < 0.001; Fig. 5d). In both cases, the surface area of food available was a better predictor of the impact of adult crowding than total air volume.

**Figure 5:**
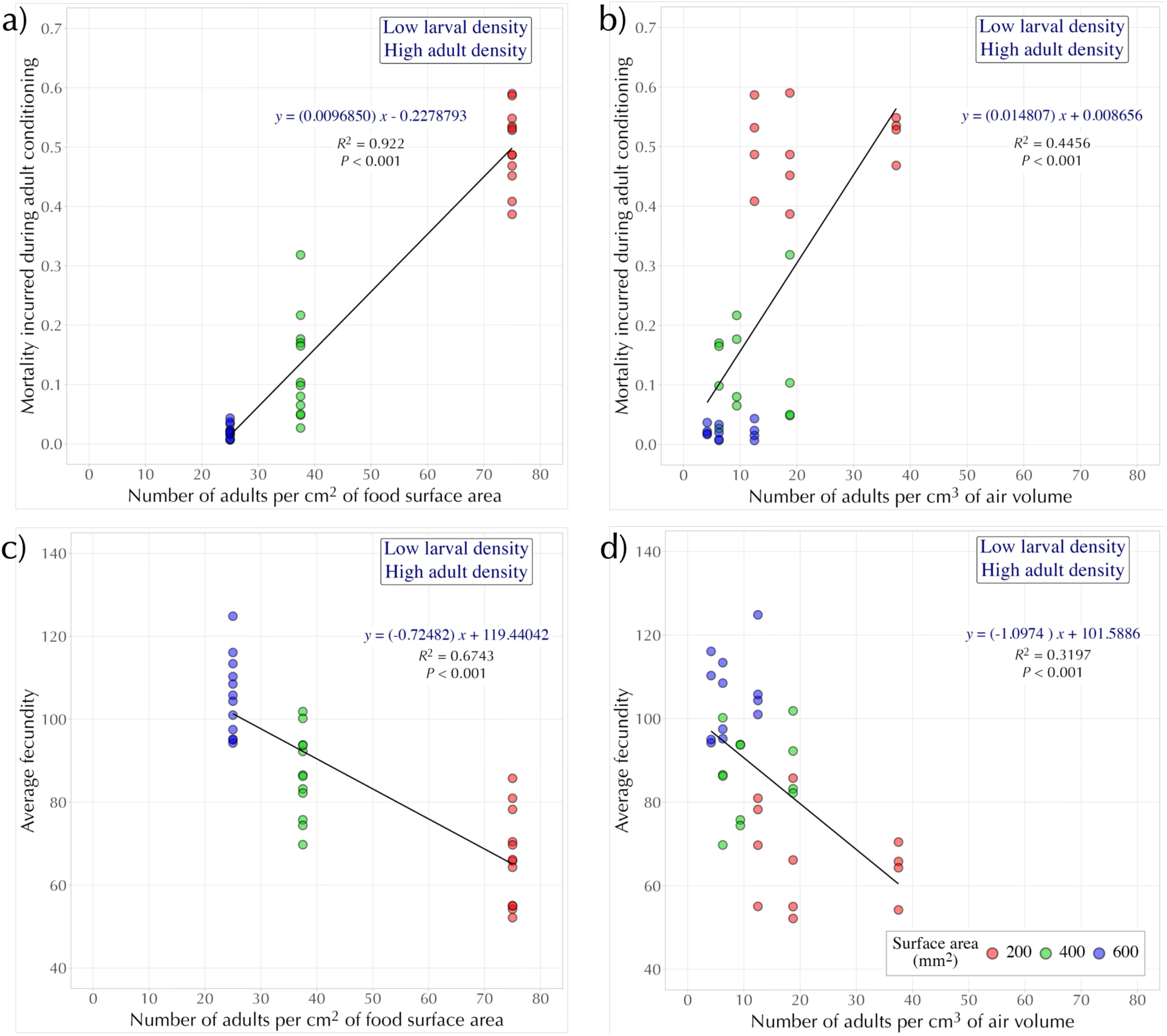
Linear least-squares regressions for flies reared at L-LD and exposed to H-AD. **(a,b)** Mean mortality incurred during adult conditioning and **(c,d)** mean female fecundity after adult conditioning as response variables. **a)** and **c)** represent regressions using the number of adults per cm^2^ of food surface area as a predictor variable, while **b)** and **d)** represent regressions using the number of adults per cm^3^ of air volume as a predictor variable.

The regression analysis established that the surface area of food available played a significant role in determining the impact of adult crowding. However, this applied only to large flies at H-AD. One possible explanation could be that the crowding intensity was much lower for small flies than that experienced by large flies. To validate this speculation, we calculated the *total biomass/surface area* ratio (henceforth *biomass/SA*; calculated as the average weight of a fly × the number of adults per unit surface area) for each combination of the vial diameter, larval and adult density. The average dry weight measurements of the flies just before experiencing adult conditioning were used to calculate the ratios. As the ratio considered the average weight of the flies and the number of individuals present in the vial per unit surface area of food available, it can be regarded as a composite measure of the intensity of crowding experienced by the flies.

The ratios obtained across the experiment are tabulated in Table 1. At L-AD, the ratios of *biomass/SA* are noticeably low for both large and small flies, indicating low stress levels. At H-AD, small flies experienced the highest intensity of crowding in the narrow vials (*biomass/SA* = 0.088). Incidentally, this is intermediate to the intensity experienced by large flies in wide and regular vials, respectively (*biomass/SA* = 0.068 and 0.103, respectively). As the stress levels across the three treatment combinations are comparable, we expected to observe a similar net effect on fecundity, regardless of the size of the flies. Consistent with our expectations, fecundity increased with adult density in all the above-mentioned treatments. The only net negative effect on fecundity observed in our experiment was for the large flies housed in the narrow vials at H-AD (*biomass/SA* = 0.206). For the small flies, no treatment had a comparable high-stress intensity and, therefore, might have contributed to the lack of adverse effects of adult crowding on fecundity.

**Table 1:** Ratios of total biomass per unit surface area for all treatment combinations. Combinations with approximately similar ratios in each category are highlighted with the same colour.

| Vial type | Surface area (mm<sup>2</sup>) | Height of air column (mm) | Volume (mm<sup>3</sup>) | Average weight of a large fly (mg) | Average weight of a small fly (mg) | Number of adults | *(Total biomass)/SA*<br>*(Avg. weight × no. of adults)*<br>Food surface area |  |
| --- | --- | --- | --- | --- | --- | --- | --- | --- |
|  |  |  |  |  |  |  | Large flies | Small flies |
| Narrow | 200 | 20 | 4000 | 0.275 | 0.118 | 150 | 0.2061 | 0.0884 |
|  |  | 40 | 8000 |  |  | 150 | 0.2061 | 0.0884 |
|  |  | 60 | 12000 |  |  | 150 | 0.2061 | 0.0884 |
| Regular | 400 | 20 | 8000 |  |  | 150 | 0.1030 | 0.0442 |
|  |  | 40 | 16000 |  |  | 150 | 0.1030 | 0.0442 |
|  |  | 60 | 24000 |  |  | 150 | 0.1030 | 0.0442 |
| Wide | 600 | 20 | 12000 |  |  | 150 | 0.0687 | 0.0294 |
|  |  | 40 | 24000 |  |  | 150 | 0.0687 | 0.0294 |
|  |  | 60 | 36000 |  |  | 150 | 0.0687 | 0.0294 |
| Narrow | 200 | 20 | 4000 |  |  | 10 | 0.0137 | 0.0059 |
|  |  | 40 | 8000 |  |  | 10 | 0.0137 | 0.0059 |
|  |  | 60 | 12000 |  |  | 10 | 0.0137 | 0.0059 |
| Regular | 400 | 20 | 8000 |  |  | 10 | 0.0068 | 0.0029 |
|  |  | 40 | 16000 |  |  | 10 | 0.0068 | 0.0029 |
|  |  | 60 | 24000 |  |  | 10 | 0.0068 | 0.0029 |
| Wide | 600 | 20 | 12000 |  |  | 10 | 0.0045 | 0.0019 |
|  |  | 40 | 24000 |  |  | 10 | 0.0045 | 0.0019 |
|  |  | 60 | 36000 |  |  | 10 | 0.0045 | 0.0019 |

## Discussion

In an earlier study (Rao *et al*., 2025) we challenged the prevailing notion that adult crowding results in adverse effects on mortality and fecundity and showed that these effects were dependent on the body size of the flies. Small body sizes obtained by the imposition of larval crowding or long-term selection for rapid development aided the flies in dealing with the adult crowding better, as indicated by an increase in fecundity with an increase in adult density, with little to no change in mortality (Rao *et al*., 2025). Further, we hypothesized a hump-shaped relationship between female fecundity and adult density – an increase in number of individuals in the container would produce a positive impact on fecundity until a threshold (positive effects zone), beyond which it would result in a net negative impact (negative effects zone). In the present study, we explored how the effects of adult crowding are shaped by altering the air volume available to the adults alongside body size. The air volume was altered via changes in the vial diameter or the height of the air column.

By altering the air volume available to the flies, we expected to modify the intensity of stress experienced by the flies during an episode of adult crowding, thereby inducing either positive or negative effects of being crowded. In our experimental design, the narrow vials would create the most stressful environment, followed by the regular and wide vials respectively. Similarly, the 20 mm air column height was expected to induce the highest level of stress, followed by the 40 mm and 60 mm heights, respectively. Additionally, large flies (reared at L-LD) would experience more stress as the effective adult density in the vials is higher than that of small flies (reared at H-LD) under similar conditions. We expected the more stressful conditions to result in higher mortality during the conditioning period and lower female fecundity subsequently.

Alterations to the air column height had negligible effects on fecundity and mortality (Fig. 3; Table S1). The intensity of adult crowding was primarily affected by the vial diameter, which, in turn, determines the surface area of food available to the adults (Figs. 2 & 3). In the vials, crowding occurs multiple times near the food surface as the flies acquire nutrition or oviposit. Females are more likely to regularly visit the food surface to oviposit or feed due to their higher nutritional needs, leading to higher female mortality during adult crowding. A decrease in the vial diameter resulted in significantly lower fecundity and higher mortality when flies were subjected to H-AD. Moreover, these effects were limited to the large flies (Fig. 2).

Our results are consistent with previous studies (Pearl & Parker, 1922; Pearl, 1932; Robertson & Sang, 1944) that emphasized the role of surface area in shaping female fecundity. In these studies, fecundity was measured during the adult crowding period, implying that the observed effects on fecundity could either arise due to differences in surface area available for oviposition or due to females being subjected to varying degrees of stress across treatments. In contrast, our assay measured fecundity after the adult conditioning period and females were housed singly during the oviposition window, eliminating any confounding effects of the presence of other individuals. Moreover, these flies encountered different vial types only during the adult conditioning period and were subsequently provided with standardized 8-dram vials (2.2 cm dia × 9.6 cm ht) for oviposition. The similarity in the results across these studies therefore suggests that the positive effects of an increase in surface area on fecundity extend beyond immediate access to a greater area for oviposition during the adult crowding episode.

### Size dependent effects

Large flies showed a context-dependent response to adult crowding based on the vial diameter – as adult density increased, mortality increased in narrow and regular vials but did not change in wide vials, while fecundity increased in regular and wide vials but decreased in narrow vials. Surprisingly, small flies showed a consistent response of an increase in fecundity with an increase in adult density that was coupled with little to no change in mortality regardless of the vial diameter (Fig. 2). Subsequently, regression analyses revealed that surface area of food plays a more prominent role than air volume in shaping the responses to crowding, albeit only for the large flies at H-AD (Fig. 5). These results indicate that body size plays a significant role in shaping the responses to adult crowding. Based on our hypothesis of a hump-shaped relationship between adult density and fecundity (Rao *et al*., 2025), we successfully explored the zones of positive and negative effects for the large flies in the present study.

In comparison, only positive effects were observed for the small flies. One possibility is that the small flies did not experience space limitation to the extent that the large flies did. For the small flies, the highest stress conditions corresponded to being housed in the narrow vials, and the *biomass/SA* ratio of this treatment is intermediate to the ratios obtained for the large flies housed in the regular and wide vials, which supports our hypothesis of small flies experiencing less stress in our experiment (Table 1). Another interesting finding is that for the small flies, the fecundity increases to the same degree across all vial types as density increases. In *Drosophila*, female fecundity strongly correlates to body size (Robertson, 1957; Lefranc & Bundgaard, 2000; Byrne & Rice, 2006), and being small could constrain egg production, thereby posing an upper limit on the extent to which fecundity can increase. While small flies may be experiencing less stress than large ones, their smaller size may also pose an upper limit for how much fecundity can increase. Thus, even if the positive effects of crowding outweigh the harmful effects, it might not result in further increased fecundity.

Incidentally, the response of large flies observed here differs from that of the JBs (reared at L-LD) from our earlier study, where fecundity decreased with increased adult density (Rao *et al*., 2025). The regular vial with the plug inserted at a height of 60 mm from the food surface corresponds to the dimensions of the vials used in the earlier study. In the present study, flies subjected to these treatment conditions showed an increase in fecundity with adult density. One plausible reason for this discrepancy is the presence of a male during the oviposition window in the earlier study. Males are known to have a detrimental effect on females with prolonged exposure – through increased mate harassment or copulation (Partridge *et al*., 1987; Fowler & Partridge, 1989; Chapman *et al*., 1995), which could be one of the reasons for the decrease in fecundity in our earlier report but not in the present study, as females were housed alone during the oviposition window. While there is some evidence that male presence during the oviposition window does not significantly impact female fecundity (Turner & Anderson, 1983; R.S. Bindya & A. Joshi, unpublished pilot data), assay-to-assay variation could also result in the observed differences in fecundity patterns. These differences, however, do not alter our conclusions regarding the role of body size and the intensity of stress experienced across differing vial types.

### Is the biomass/SA ratio a reliable index of the crowding intensity?

The *biomass/SA* ratio can be altered based on three factors – the food surface area, the number of adults and their body weight. We hypothesize that the *biomass/SA* ratio reflects the *effective adult density* during crowding and can be considered a good index of the intensity of adult crowding. The ratios were used to compare the stress likely experienced across vial types for flies of different sizes (Table 1). For treatments with a ratio < 0.10, very low mortality levels were observed (< 15%), regardless of the size of the adults (Fig. 6a). While the range of ratios in this zone (values < 0.10) differed up to 45-fold, variation in mortality was minimal (< 10%). Our experiment had only two treatment combinations with a *biomass/SA* ratio > 0.1, with the difference between them being only twofold. However, the mortality was ∼30% higher in the treatment with the higher *biomass/SA* ratio, suggesting that ratios above 0.1 could result in flies experiencing a significantly higher amount of stress.

**Figure 6:**
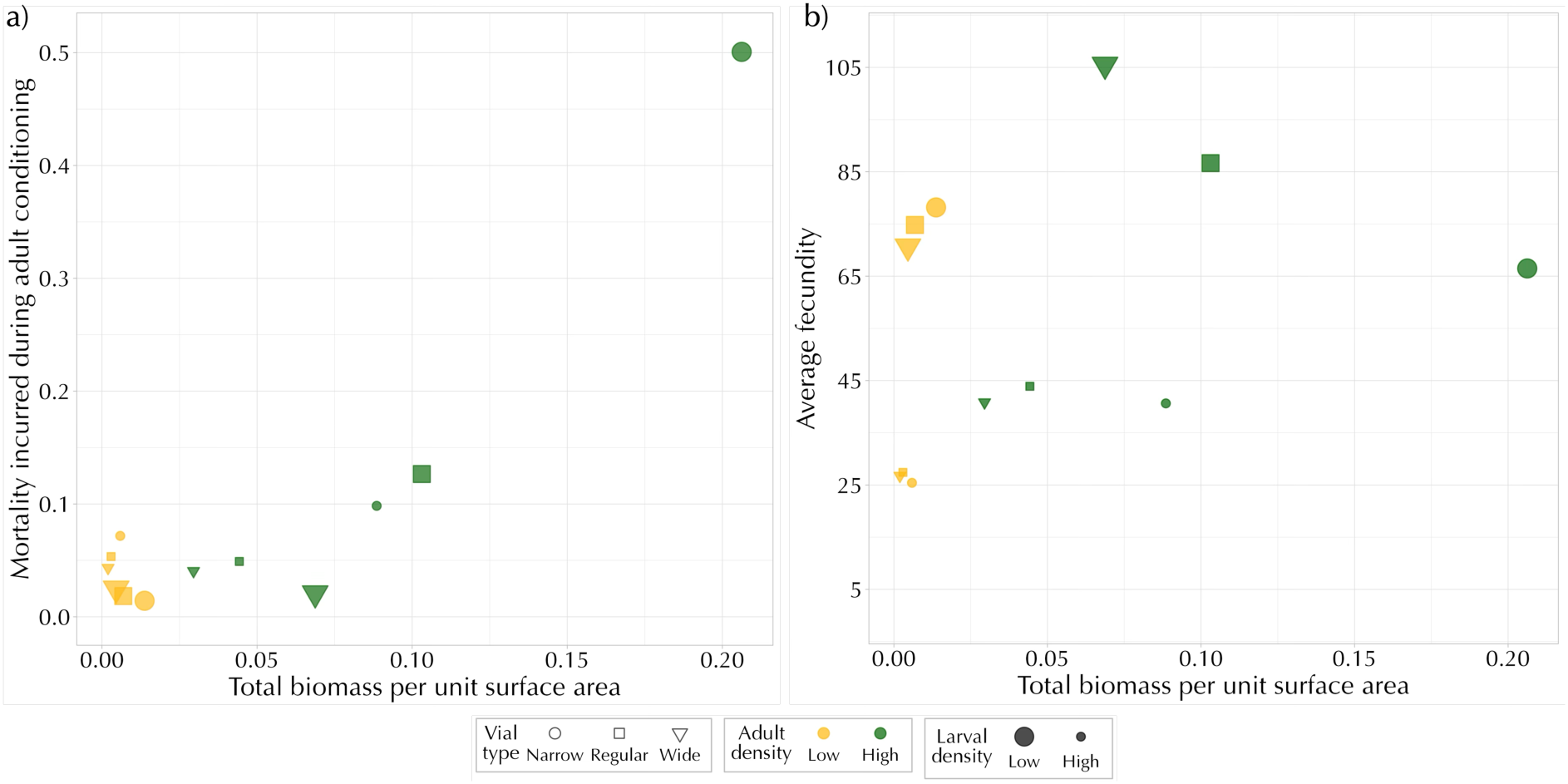
a) Mean mortality and **b)** Mean fecundity corresponding to the *total biomass per unit surface area* ratios obtained across all treatment combinations.

Fecundity, on the other hand, seems to be primarily determined by the size of the flies. In any given treatment combination, large flies consistently experienced ∼2.34 times the stress (*biomass/SA*) experienced by small flies (Table 1). Nevertheless, the absolute fecundity of large flies was always higher, indicating that the size of flies probably played a more prominent role than the *biomass/SA* in determining fecundity. An increase in adult density results in a concomitant *biomass/SA* ratio increase. Under these conditions, small flies consistently responded with increased fecundity to an increased *biomass/SA* ratio despite the stress experienced in vials increasing threefold from the wide to the narrow vial treatment combinations (Fig. 6b, Table 1). In contrast, large flies showed increased fecundity at H-AD in the wide and regular vials, and decreased fecundity was observed only in the narrow vials. Interestingly, regardless of the size of the flies, an increase in the *biomass/SA* ratio (via an increase in adult numbers) led to a concomitant rise in fecundity when the ratio remained ⪅ 0.1. A decrease in fecundity was observed when the ratio was much higher than 0.1. Our experiment, incidentally, only explored this range of values for the large flies. It suggests one of the following two possibilities:

1. Flies have body-size-dependent reaction norms to changes in the *biomass/SA* ratio brought about via changes in the number of adults or the surface area of food available.
2. Flies, irrespective of body size, have similar reaction norms, and a common threshold of *biomass/SA* exists, beyond which flies are negatively affected. Our study’s range of ratios can offer limited speculation about this possibility (as only two treatments had values > 0.1). Future studies could explore more treatment combinations with *biomass/SA* ≳ 0.1 for flies of both sizes to validate this speculation.

Given the observed data for the small flies, the threshold proposed above could hold, or small flies could have a different response norm relative to the large flies. The proposed models - a hump-shaped relationship between fecundity and adult density (Rao *et al*., 2025) or the *biomass/SA* threshold – do not entirely explain the consistent increase in fecundity observed in the small flies. Based on the model proposed from our earlier work, we expected that even if fecundity increased at H-AD, the extent of the increase in fecundity in each treatment would be different owing to varying levels of stress experienced. However, fecundity increased by approximately the same amount across all vial types with increased adult density. While this aspect can be attributed to the body-size-dependent constraints detailed in the previous sub-section, it would not correspond well with a hump-shaped relationship between fecundity and adult density. Furthermore, our study only explored a limited range of *biomass/SA* values, with only one treatment combination - large flies, housed in narrow vials with an air column height of 20 mm (*biomass/SA* = 0.2) - corresponding to a net negative effect when crowding was experienced, limiting the generalization of these results (Figs. 3 & 6). Therefore, further studies are required to test which of the two models better explains the observed results for small flies.

### Future directions

In the present study, small flies possibly did not experience space limitation to the same extent as large flies (Table 1). Examining treatments with a higher range of *biomass/SA* ratios for the small flies would be worthwhile in assessing the validity of the hump-shaped relationship between fecundity and density that was postulated in our earlier study (Rao *et al*., 2025). It could be done with vials of even smaller diameter than in the present study or by subjecting small flies to higher intensities of crowding. Furthermore, it would be interesting to investigate if changes of similar magnitudes brought about by altering either surface area, body size, or number of individuals (one at a time) could result in similar effects on mortality and fecundity after experiencing a short duration of adult crowding.

Small flies in our study were obtained by imposing larval crowding. One might speculate if the observed results were solely due to body size differences or if the process of experiencing larval crowding itself, over and above the body size, could have shaped the response. Future studies could explore the response of small flies achieved through other means (reviewed in Shingleton *et al*., 2009), such as rearing at high temperatures (Cavicchi *et al*., 1989; Partridge *et al*., 1994; Nunney & Cheung, 1997), direct selection for smaller body sizes (Partridge & Fowler, 1993; Partridge *et al*., 1999) and nutrient limitation (Robertson, 1963). In this manner, it might be possible to differentiate the roles of body size and larval crowding in shaping the response to adult crowding.

Studies have shown that among sets of populations adapted to larval crowding (CUs), adult crowding (UCs) and their controls (UUs), CUs fared the worst under conditions of adult crowding, suggesting that adaptation to larval crowding might trade-off with adaptations to adult crowding (Joshi A. 1997; Joshi *et al*. 1998). In those studies, adult crowding was imposed on large flies, as all populations were reared at low larval density for the assay (∼60-70 eggs in ∼6 mL of food). This larval density represents the native larval density regime usually experienced by the UUs and UCs. CUs, on the other hand, experience crowding as larvae (∼1200 eggs in ∼6 mL of food) and were tested in the assay under non-native larval rearing conditions. Therefore, the effects observed in the study could result from the interactive effects of selection regime and body size (and/or larval rearing density). It might be worthwhile to subject flies from such selection regimes to a range of larval densities (including native densities) and then examine their response to adult crowding using a full-factorial design.

Overall, our findings indicate that the relationship between the intensity of stress experienced and the space available in the container is quite nuanced, as has also been found to be the case for intensity of larval crowding and food volume (Venkitachalam *et al*., 2023). If one assumes that crowding is primarily experienced near the surface of the food, changing the space available by altering the air column height would not substantially affect the crowding intensity. At the same time, alterations to the vial diameter would have a significant effect. Our results emphasize the nuanced effects of adult crowding and highlight the importance of considering the body size of the flies subjected to crowding and the food surface area provided to them.

## Acknowledgements

We thank Bhawna Mittal, Mohnish Singh, Rajanna N. and Muniraju P. for their assistance in the laboratory. MR was supported by a doctoral fellowship from JNCASR. The study was supported by intramural funds from JNCASR and, in part, by A. Joshi’s personal funds.

## Author contributions

MR, CT and AJ conceptualized this study; MR, CT, BRS and AJ designed the experiment; MR, CT and BRS carried out the experiments; MR analyzed the data and wrote the first draft of the paper; MR, CT and AJ contributed to subsequent revisions and the final draft of the paper.

## References

Borash, D. J., Gibbs, A. G., Joshi, A., & Mueller, L. D. (1998). A genetic polymorphism maintained by natural selection in a temporally varying environment. The American Naturalist, 151(2), 148–156. 10.1086/286108

Borash, D. J., & Ho, G. T. (2001). Patterns of selection: Stress resistance and energy storage in density-dependent populations of *Drosophila melanogaster*. Journal of Insect Physiology, 47(12), 1349–1356. 10.1016/S0022-1910(01)00108-1

Byrne, P. G., & Rice, W. R. (2006). Evidence for adaptive male mate choice in the fruit fly *Drosophila melanogaster*. Proceedings of the Royal Society B: Biological Sciences, 273(1589), 917–922. 10.1098/rspb.2005.3372

Cavicchi, S., Guerra, D., Natali, V., Pezzoli, C., & Giorgi, G. (1989). Temperature-related divergence in experimental populations of *Drosophila melanogaster*. II. Correlation between fitness and body dimensions. Journal of Evolutionary Biology, 2(4), 235–251. 10.1046/j.1420-9101.1989.2040235.x

Chapman, T., Liddle, L. F., Kalb, J. M., Wolfner, M. F., & Partridge, L. (1995). Cost of mating in *Drosophila melanogaster* females is mediated by male accessory gland products. Nature, 373(6511), 241–244. 10.1038/373241a0

Chapman, T., & Partridge, L. (1997). Female fitness in *Drosophila melanogaster*: An interaction between the effect of nutrition and of encounter rate with males. Proceedings of the Royal Society of London. Series B: Biological Sciences, 263(1371), 755–759. 10.1098/rspb.1996.0113

Fowler, K., & Partridge, L. (1989). A cost of mating in female fruitflies. Nature, 338(6218), 760– 761. 10.1038/338760a0

Joshi, A. (1997). Laboratory studies of density-dependent selection: Adaptations to crowding in *Drosophila melanogaster*. Current Science, 72(8), 555–562.

Joshi, A., & Mueller, L. D. (1988). Evolution of higher feeding rate in *Drosophila* due to density-dependent natural selection. Evolution, 42(5), 1090–1093. 10.2307/2408924

Joshi, A., & Mueller, L. D. (1993). Directional and stabilizing density-dependent natural selection for pupation height in *Drosophila melanogaster*. Evolution, 47(1), 176–184. 10.1111/j.1558-5646.1993.tb01208.x

Joshi, A., & Mueller, L. D. (1996). Density-dependent natural selection in *Drosophila*: Trade-offs between larval food acquisition and utilization. Evolutionary Ecology, 10(5), 463–474. 10.1007/BF01237879

Joshi, A., Wu, W.-P., & Mueller, L. D. (1998). Density-dependent natural selection in *Drosophila*: Adaptation to adult crowding. Evolutionary Ecology, 12(3), 363–376. 10.1023/A:1006508418493

Lefranc, A., & Bundgaard, J. (2000). The influence of male and female body size on copulation duration and fecundity in *Drosophila melanogaster*. Hereditas, 132(3), 243–247. 10.1111/j.1601-5223.2000.00243.x

Mueller, L. D. (1997). Theoretical and empirical examination of density-dependent selection. Annual Review of Ecology, Evolution, and Systematics, 28, 269–288. 10.1146/annurev.ecolsys.28.1.269

Nagarajan, A., Natarajan, S. B., Jayaram, M., Thammanna, A., Chari, S., Bose, J., Jois, S. V., & Joshi, A. (2016). Adaptation to larval crowding in *Drosophila ananassae* and *Drosophila nasuta nasuta*: Increased larval competitive ability without increased larval feeding rate. Journal of Genetics, 95(2), 411–425. 10.1007/s12041-016-0655-9

Nunney, L., & Cheung, W. (1997). The effect of temperature on body size and fecundity in female *Drosophila melanogaster*: Evidence for adaptive plasticity. Evolution, 51(5), 1529–1535. 10.1111/j.1558-5646.1997.tb01476.x

Partridge, L., & Fowler, K. (1993). Responses and correlated responses to artificial selection on thorax length in *Drosophila melanogaster*. Evolution, 47(1), 213–226. 10.1111/j.1558-5646.1993.tb01211.x

Partridge, L., Green, A., & Fowler, K. (1987). Effects of egg-production and of exposure to males on female survival in *Drosophila melanogaster*. Journal of Insect Physiology, 33(10), 745–749. 10.1016/0022-1910(87)90060-6

Partridge, L., Langelan, R., Fowler, K., Zwaan, B., & French, V. (1999). Correlated responses to selection on body size in *Drosophila melanogaster*. Genetics Research, 74(1), 43–54. 10.1017/S0016672399003778

Pearl, R. (1932). The influence of density of population upon egg production in *Drosophila melanogaster*. Journal of Experimental Zoology, 63(1), 57–84. 10.1002/jez.1400630103

Pearl, R., Miner, J. R., & Parker, S. L. (1927). Experimental studies on the duration of life. XI. Density of population and life duration in *Drosophila*. The American Naturalist, 61(675), 289–318. 10.1086/280154

Pearl, R., & Parker, S. L. (1922). On the influence of density of population upon the rate of reproduction in *Drosophila*. Proceedings of the National Academy of Sciences, 8(7), 212–219. 10.1073/pnas.8.7.212

Prasad, N. G., & Joshi, A. (2003). What have two decades of laboratory life-history evolution studies on *Drosophila melanogaster* taught us? Journal of Genetics, 82(1), 45–76. 10.1007/BF02715881

R Core Team (2024). R: A Language and Environment for Statistical Computing. R Foundation for Statistical Computing, Vienna, Austria. https://www.r-project.org/

Rao, M., Temura, C., Mital, A., Anvitha, S., & Joshi, A. (2025). Bigger is not always better: Size-dependent fitness effects of adult crowding in *Drosophila melanogaster*. bioRxiv, 2025.04.21.649761. 10.1101/2025.04.21.649761

Robertson, F. W. (1957). Studies in quantitative inheritance XI. Genetic and environmental correlation between body size and egg production in *Drosophila melanogaster*. Journal of Genetics, 55(3), 428–443. 10.1007/BF02984061

Robertson, F. W. (1963). The ecological genetics of growth in *Drosophila* 6. The genetic correlation between the duration of the larval period and body size in relation to larval diet. Genetics Research, 4(1), 74–92. 10.1017/S001667230000344X

Robertson, F. W., & Sang, J. H. (1944). The ecological determinants of population growth in a *Drosophila* culture. I. Fecundity of adult flies. Proceedings of the Royal Society of London. Series B - Biological Sciences, 258–277. 10.1098/rspb.1944.0017

Sarangi, M. (2018). *Ecological details mediate different paths to the evolution of larval competitive ability in* Drosophila [Thesis, Jawaharlal Nehru Centre for Advanced Scientific Research]. https://libjncir.jncasr.ac.in/xmlui/handle/10572/2722

Sarangi, M., Nagarajan, A., Dey, S., Bose, J., & Joshi, A. (2016). Evolution of increased larval competitive ability in *Drosophila melanogaster* without increased larval feeding rate. Journal of Genetics, 95(3), 491–503. 10.1007/s12041-016-0656-8

Sheeba, V., Madhyastha, N. A. A., & Joshi, A. (1998). Oviposition preference for novel versus normal food resources in laboratory populations of *Drosophila melanogaster*. Journal of Biosciences, 23(2), 93–100. 10.1007/BF02703000

Shingleton, A. W., Estep, C. M., Driscoll, M. V., & Dworkin, I. (2009). Many ways to be small: Different environmental regulators of size generate distinct scaling relationships in *Drosophila melanogaster*. Proceedings of the Royal Society B: Biological Sciences, 276(1667), 2625–2633. 10.1098/rspb.2008.1796

Shiotsugu, J., Leroi, A. M., Yashiro, H., Rose, M. R., & Mueller, L. D. (1997). The symmetry of correlated selection responses in adaptive evolution: An experimental study using *Drosophila*. Evolution, 51(1), 163–172. 10.1111/j.1558-5646.1997.tb02397.x

Turner, M. E., & Anderson, W. W. (1983). Multiple mating and female fitness in *Drosophila pseudoobscura*. Evolution, 37(4), 714–723. 10.2307/2407913

Venkitachalam, S. (2023). The ecology and evolution of larval competitive ability in laboratory populations of *Drosophila* [Thesis, Jawaharlal Nehru Centre for Advanced Scientific Research (JNCASR)]. https://libjncir.jncasr.ac.in/xmlui/handle/123456789/3427

Venkitachalam, S., Sajith, V. S., & Joshi, A. (2023). More than just density: The role of egg number, food volume and container dimensions in mediating larval competition in *Drosophila melanogaster* (p. 2023.07.26.550621). bioRxiv. 10.1101/2023.07.26.550621

Wickham, H. (2016). ggplot2: Elegant graphics for data analysis. Springer-Verlag. 10.1007/978-3-319-24277-4

Wickham, H., Averick, M., Bryan, J., Chang, W., McGowan, L. D., François, R., Grolemund, G., Hayes, A., Henry, L., Hester, J., Kuhn, M., Pedersen, T. L., Miller, E., Bache, S. M., Müller, K., Ooms, J., Robinson, D., Seidel, D. P., Spinu, V., … Yutani, H. (2019). Welcome to the Tidyverse. Journal of Open Source Software, 4(43), 1686. 10.21105/joss.01686

