## Supplementary Materials for "Fitness effects of adult crowding in *Drosophila*: more than just overall density"

#### Supplementary tables

**Table S1:** ANOVA results for the effect of vial diameter (surface area of food), height of air column, larval and adult density on adult mortality during two days of adult conditioning. In this design, random factors and their interactions are not tested for significance and are omitted from the table.

| **Effect** | ***df*** | **MS** | ***F*** | ***P*** |
|---|---|---|---|---|
| Vial diameter (Surface area) | 2 | 0.515 | 153.895 | < 0.001 |
| Larval density | 1 | 0.246 | 19.060 | 0.022 |
| Adult density | 1 | 0.741 | 75.965 | 0.003 |
| Height | 2 | 0.002 | 1.607 | 0.275 |
| Sex | 1 | 0.353 | 83.243 | 0.002 |
| Surface area × Larval density | 2 | 0.243 | 65.103 | < 0.001 |
| Surface area × Adult density | 2 | 0.450 | 135.166 | < 0.001 |
| Larval density × Adult density | 1 | 0.652 | 101.744 | 0.002 |
| Surface area × Height | 4 | 0.002 | 1.040 | 0.426 |
| Larval density × Height | 2 | 0.001 | 0.907 | 0.452 |
| Adult density × Height | 2 | 0.002 | 2.452 | 0.166 |
| Surface area × Sex | 2 | 0.188 | 34.328 | < 0.001 |
| Larval density × Sex | 1 | 0.404 | 437.987 | < 0.001 |
| Adult density × Sex | 1 | 0.513 | 512.945 | < 0.001 |
| Height × Sex | 2 | 0.005 | 2.479 | 0.164 |
| Surface area × Larval density × Adult density | 2 | 0.346 | 110.164 | < 0.001 |
| Surface area × Larval density × Height | 4 | 0.002 | 2.254 | 0.123 |
| Surface area × Adult density × Height | 4 | 0.002 | 1.613 | 0.234 |
| Larval density × Adult density × Height | 2 | 0.0002 | 0.098 | 0.907 |
| Surface area × Larval density × Sex | 2 | 0.222 | 69.687 | < 0.001 |
| Surface area × Adult density × Sex | 2 | 0.230 | 35.568 | < 0.001 |
| Larval density × Adult density × Sex | 1 | 0.352 | 157.341 | 0.001 |
| Surface area × Height × Sex | 4 | 0.0003 | 0.272 | 0.890 |
| Larval density × Height × Sex | 2 | 0.0002 | 0.170 | 0.846 |
| Adult density × Height × Sex | 2 | 0.0005 | 1.157 | 0.375 |
| Surface area × Larval density × Adult density × Height | 4 | 0.0006 | 0.260 | 0.897 |
| Surface area × Larval density × Adult density × Sex | 2 | 0.173 | 59.732 | < 0.001 |

| | | | | |
|---|---|---|---|---|
| Surface area × Larval density × Height × Sex | 4 | 0.0003 | 0.140 | 0.963 |
| Surface area × Adult density × Height × Sex | 4 | 0.002 | 1.172 | 0.370 |
| Larval density × Adult density × Height × Sex | 2 | 0.0002 | 0.145 | 0.867 |
| Surface area × Larval density × Adult density × Height × Sex | 4 | 0.0006 | 1.027 | 0.432 |

**Table S2:** ANOVA results for the effect of vial diameter (food surface area), height of air column, larval and adult density on female fecundity after two days of adult conditioning. In this design, random factors and their interactions are not tested for significance and are omitted from the table.

| **Effect** | ***df*** | **MS** | ***F*** | ***P*** |
|---|---|---|---|---|
| Vial diameter (Surface area) | 2 | 852.695 | 5.295 | 0.047 |
| Larval density | 1 | 76955.356 | 92.923 | 0.002 |
| Adult density | 1 | 6527.285 | 60.973 | 0.004 |
| Height | 2 | 84.449 | 5.743 | 0.040 |
| Surface area × Larval density | 2 | 687.308 | 3.416 | 0.102 |
| Surface area × Adult density | 2 | 1548.704 | 42.435 | < 0.001 |
| Larval density × Adult density | 1 | 116.821 | 1.627 | 0.291 |
| Surface area × Height | 4 | 25.543 | 0.827 | 0.532 |
| Larval density × Height | 2 | 1.373 | 0.013 | 0.986 |
| Adult density × Height | 2 | 19.153 | 0.862 | 0.468 |
| Surface area × Larval density × Adult density | 2 | 1707.150 | 19.405 | < 0.001 |
| Surface area × Larval density × Height | 4 | 29.338 | 1.583 | 0.241 |
| Surface area × Adult density × Height | 4 | 62.268 | 4.907 | < 0.001 |
| Larval density × Adult density × Height | 2 | 12.056 | 1.539 | 0.288 |
| Surface area × Larval density × Adult density × Height | 4 | 29.877 | 1.0140 | 0.438 |

**Table S3:** ANOVA results for the effect of larval rearing density and sex on dry weight of flies. The dry weight of flies in each treatment combination represents their weight just before they experienced the adult conditioning treatments. In this design, random factors and their interactions are not tested for significance and are omitted from the table.

| **Effect** | ***df*** | **MS** | ***F*** | ***P*** |
|---|---|---|---|---|
| Larval density | 1 | 0.098 | 290.82 | < 0.001 |
| Sex | 1 | 0.027 | 42.198 | 0.007 |
| Larval density × Sex | 1 | 0.003 | 24.213 | 0.016 |

**Table S4:** Results for linear regression analyses carried out for female fecundity, total mortality and female mortality as response variables, respectively. The subset of data used for the regression analyses in each case is also provided in the table. The rows highlighted in green correspond to the plots in Figure 5.

| **Response variable** | **Subset of data used for the regression analyses** | **The surface area of food as a predictor variable ($R^2$)** | **The volume of air as a predictor variable ($R^2$)** |
|---|---|---|---|
| Fecundity | N/A | 0.0075 | 0.0048 |
| Fecundity | Low larval density (L-LD) | 0.0075 | 0.0064 |
| Fecundity | High larval density (H-LD) | 0.4056 | 0.2804 |
| Fecundity | L-LD and L-AD (Low adult density) | 0.0813 | 0.0685 |
| Fecundity | L-LD and H-AD (High adult density) | 0.6743 | 0.3196 |
| Fecundity | H-LD and L-AD | 0.0219 | 0.0272 |
| Fecundity | H-LD and H-AD | 0.0060 | 0.0080 |
| Total mortality | N/A | 0.4410 | 0.3468 |
| Total mortality | L-LD | 0.8198 | 0.6057 |
| Total mortality | H-LD | 0.1565 | 0.2340 |
| Total mortality | L-LD and L-AD | 0.0634 | 0.0094 |
| Total mortality | L-LD and H-AD | 0.9219 | 0.4456 |
| Total mortality | H-LD and L-AD | 0.1849 | 0.0795 |
| Total mortality | H-LD and H-AD | 0.4295 | 0.4832 |
| Female mortality | L-LD and L-AD | 0.0407 | 0.0009 |
| Female mortality | L-LD and H-AD | 0.9327 | 0.4632 |
| Female mortality | H-LD and L-AD | 0.0114 | 0.0274 |
| Female mortality | H-LD and H-AD | 0.2923 | 0.4306 |

### Supplementary Figures

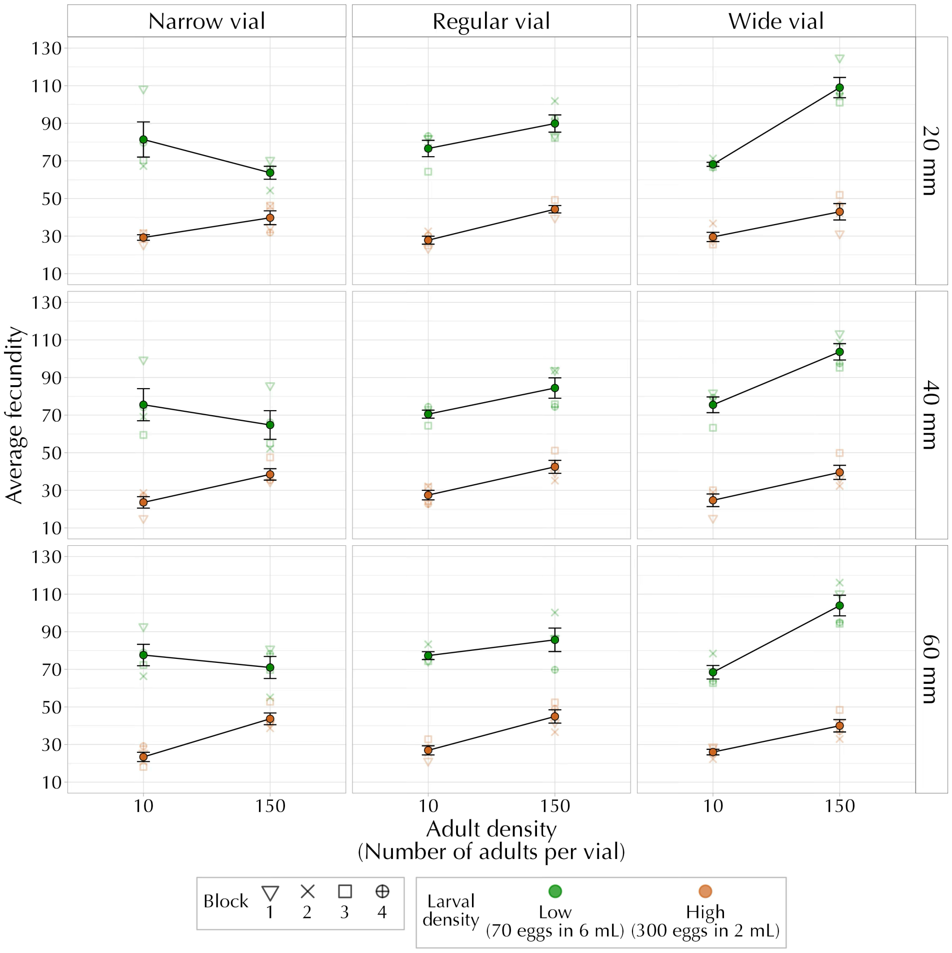


**Figure S1:** Mean female fecundity after experiencing adult conditioning for all combinations of vial diameter, air column height, larval and adult density (averaged over replicate blocks). The error bars represent the standard error around the mean.

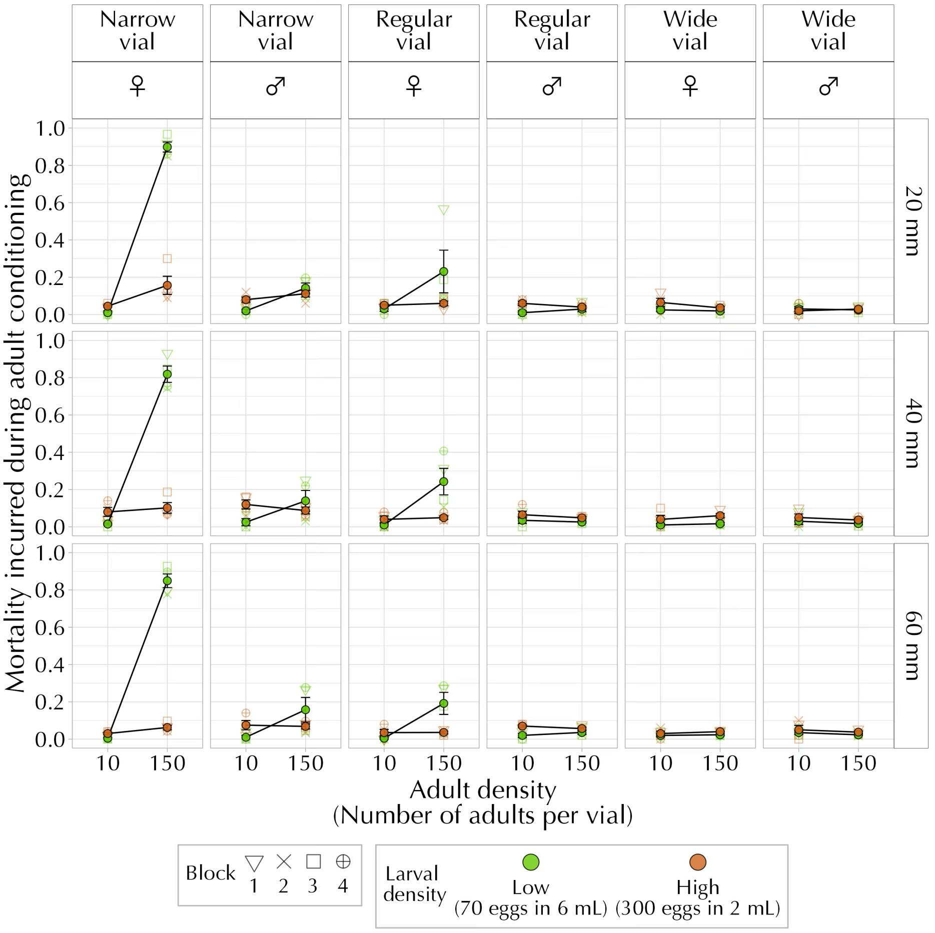


**Figure S2:** Mean mortality incurred during adult conditioning for all combinations of vial diameter, air column height, sex, larval and adult density (averaged over replicate blocks). The error bars represent the standard error around the mean.

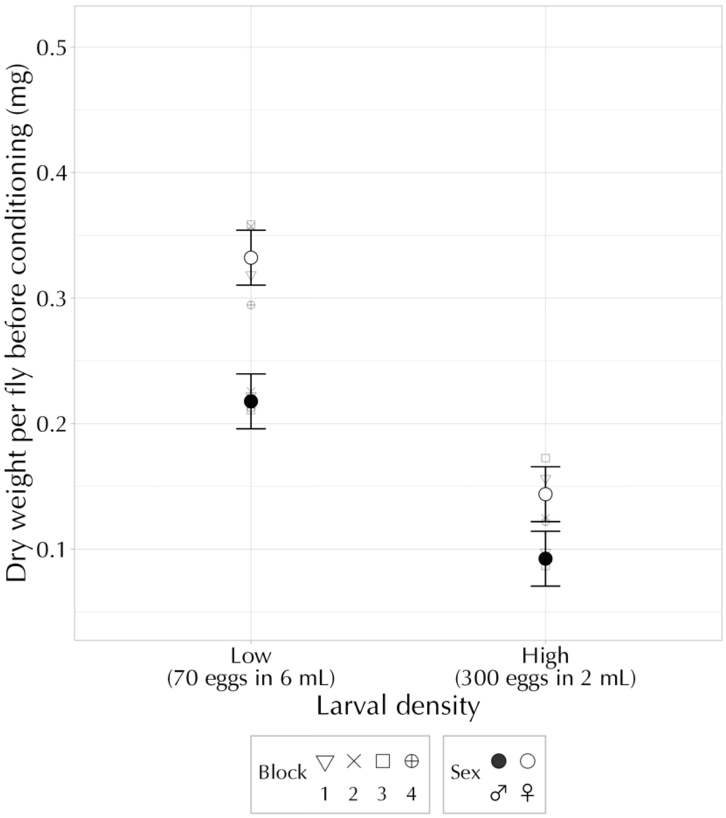


**Figure S3:** Mean dry weight per fly just before experiencing adult conditioning for all combinations of larval density and sex (averaged over replicate blocks). Error bars are 95% confidence intervals around the mean and can be used for visual hypothesis testing.
